# Prevalence of Hepatitis C Virus Genotypes in the Southern Region, Saudi Arabia

**DOI:** 10.1101/603902

**Authors:** Abdullah M AlKahtani, Meaad N Alsultan, Abdulrahim R Hakami, Mohammad Alamri

## Abstract

**Aim:** The aim of this study was to investigate the prevalence of HCV genotypes in the Southern Region, Saudi Arabia.

**Materials & methods:** A total of 76 HCV-positive individuals were selected for this study, including 34 males and 42 females, both acute and chronic patients. All HCV isolates were genotyped by direct sequencing of the 5’UTR region. The Chemistry profile, viral load and history of blood transfusion were collected from the hospital record.

**Results:** The most common genotype was gt 4 (48.7%) followed by gt 1 (34.2%) and gt 3 (14.5%). Genotype 2 (2.6%) was only found in elderly male individuals. Patients with history of blood transfusion showed a high percentage of genotype 1 compared to the total number of all patients with genotype 1 (23% and 11% respectively). Biochemical assay showed high level of ALT particularly in genotype 4. No significant relationship was observed between HCV genotype and AST level between genotypes. The viral load was higher in HCV patients received blood transfusion than other genotypes.

**Conclusion:** The prevalence of genotypes in this study confirmed the observation of other investigations, but no link was found between sex and genotype. There might be an association between blood donation in the past and infections with genotype 1.

## Introduction

Hepatitis C virus (HCV) is an enveloped, single-stranded RNA virus and belongs to Hepacivirus genus and *Flaviviridae* family [1]. HCV continues to be a worldwide critical public health concern which causes high morbidity and mortality. The virus is transmitted parenterally and uses CD8, SR-BI, claudin and occludin receptors for entry [2]. More than 170 million individuals are chronically infected with the virus worldwide with an estimation of more than 15 million seropositive individuals in the Middle East and North Africa, and this appears to be the highest prevalence with 3.6% infections [3]. It has been estimated that approximately 75%–85% of HCV-infected individuals develop chronic infection and up to half a million people die each year from hepatocellular carcinoma (HCC) and other related liver diseases [4,5].

There are about 80 recognized subtypes of HCV classified under at least six genotypes [6], and the prevalence is linked with geographical distribution [7]. The genetic diversity of the virus (quasi-species) suggests that more novel subtypes will be discovered. It has been hypothesized that HCV genotypes correlate with a varying degree of disease severity. Specific HCV genotypes were linked with liver steatosis, hepatic reactivation and liver fibrosis [8,9]. Genotype 1 is associated with higher risk of cirrhosis, HCC development, and poor response to therapy [10]. Because of that, genotyping has become necessary in the clinical analysis of patients with hepatitis C, to monitor therapeutic response and duration of treatment [11].

Despite that epidemiological data on HCV in Saudi Arabia is deficient [12,13] and also limited by small sample size [14], it has been speculated that viral hepatitis is highly endemic in the southwestern region in Saudi Arabia [15]. However, estimating the overall prevalence of HCV in the country is difficult due to lack of more studies in different geographical regions [16] with conflicting results regarding the actual prevalence [17]. It was identified that genotype 4 is the most predominant in Saudi Arabia followed by genotype 1 [18–20]. The nucleotide conservation was 92% and 95.5% respectively [21]. Subtypes 4a and 1b showed the highest level of nucleotides variation [22].

HCV is characterized by the emergence of resistance associated substitutions that renders the virus less susceptible to the direct-acting antiviral agents [23]. Better understanding of the risk factors could support the development of targeted strategies to reduce HCV infection and disease severity. For example, a lower rate of sustained virologic response (SVR) was found in patients infected with the subtype 4r more than the other subtypes [24]. More studies with large number of samples are needed to examine the role of circulating subtypes in disease severity.

A recent study suggested that genotype 4 in Saudi Arabia has declined [25,26]. In addition, data on HCV genotyping in different Saudi Arabian regions is still scarce and this necessitates further investigations. The aim of this study is to determine the prevalence of HCV genotypes in acute and chronic patients in the Southern Region of Saudi Arabia.

## Materials & methods

### • Patients selection

A total of 76 HCV-positive patients from different cities in the Southern Region (34 males and 42 females; aged: 20-79 y) seen from 2016 to 2017 at the Armed Forces Hospital Southern Region (AFHSR) were enrolled in this study. Diagnosis of chronic hepatitis C was made according to the AFHSR Guideline on the Prevention and Treatment of Hepatitis C.

### • Detection of HCV antibodies

Anti-HCV antibodies in patients’ sera were qualitatively screened using a chemiluminescent microparticle immunoassay (CMIA-ARCHITECHT3; ABBOTT). Recombinant blot immunoassay (RIBA) were performed using Line ImmunoAssay for the detection of anti-HCV antibodies in human serum which was performed according to the manufacturer instructions (INNO-LIA HCV Score). Seropositive samples were examined for liver function tests, direct bilirubin levels and processed for genotyping determination.

### • Biochemistry profile

Blood samples were collected in plain tubes and centrifuged for 10min at 3000rpm. Patients’ sera were tested for biochemistry profile according to manufacturer’s instructions using the fully automated machine (Beckman Unicel DxC 8000) in which the level of aspartate aminotransferase (AST), alanine aminotransferase (ALT), alkaline phosphatase (ALP) and direct bilirubin were determined.

### • Genotyping of HCV

For HCV genotyping and HCV-RNA quantitative determination, 10ml of EDTA-anticoagulated blood were collected and transported promptly to Bioscientia Lab (Germany) on the same day of collection. Briefly, cobas^®^ HCV GT assay (Roche Diagnostics) was run on the cobas 4800 system for HCV genotyping, which relies on direct sequencing of the NS5B gene on AB capillary sequencing system.

### • Statistical analysis

Data were analyzed using GraphPad Prism, version 7.04 (GraphPad Software, CA, USA). The results of liver function tests are presented as the mean and percentage of overall patients infected with a specific genotype.

### • Ethics statement

The project and data written were approved by the Ethics Committee of the College of Medicine, King Khalid University (KKU) and from AFHSR.

## Results

### • Chemistry profile and immunoassays

A total of 76 diagnosed HCV positive cases were confirmed by RIBA and selected for biochemistry profile for liver function tests, quantitative determination of HCV RNA and HCV genotyping. Of the 76 patients included in the study, 34 males (45%) and 42 females (55%) were all Saudi Arabian (20–79 years).

From different clinics at AFHSR, 76 patients were selected for enzyme immunoassay. Table 1 and Figure 1 show the percentage and the mean of AST and ALT, which are important markers of hepatocytes injury. The liver function tests were determined for all HCV-seropositive patients. Patients were further investigated using chemiluminescent microparticle immunoassay (CMIA) and RIBA. Thirty patients (39%) were asymptomatic and showed normal biochemistry profile, while nearly 46% of the all patients showed a significant positive relationship between different HCV genotypes (specifically genotype 4) and the abnormal level of AST and ALT. However, no relationship was found between HCV infected individuals and the level of direct bilirubin and alkaline phosphatase (data not shown). Seventy-six positive patients were further investigated for genotyping and quantitative determination of the viral load for follow up.

**Figure 1.**
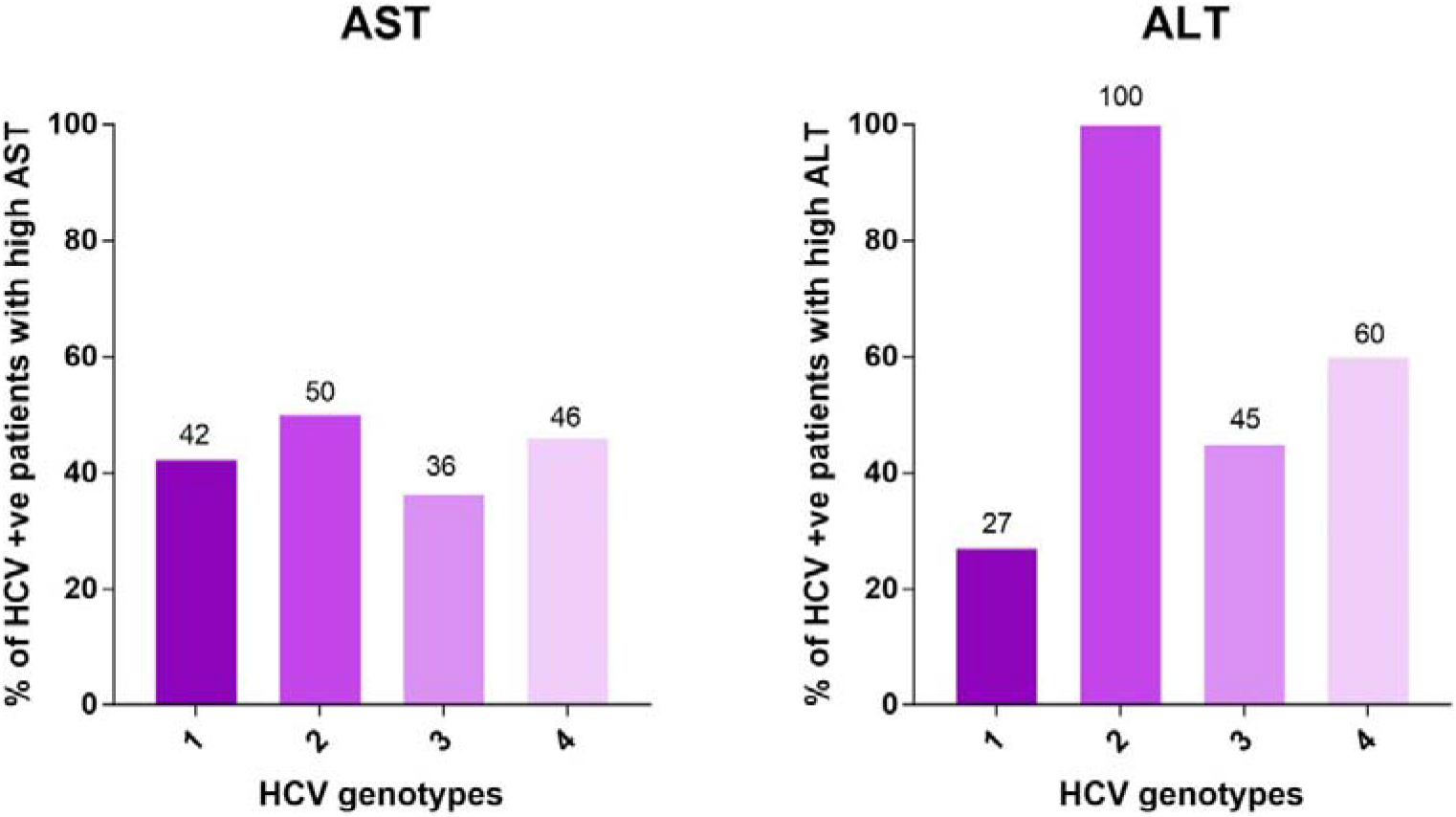
The percentage and mean of AST and ALT in all HCV genotypes determined. Figure 1 shows the level of AST and ALT versus HCV genotypes

**Table 1.**
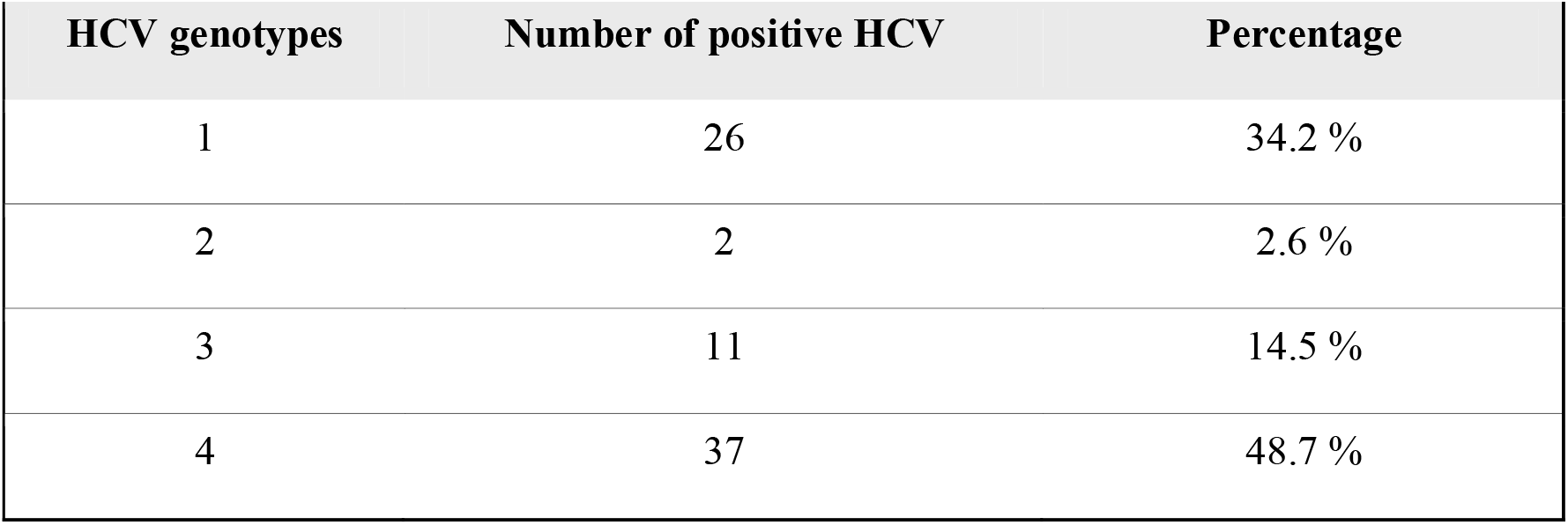
Percentage and mean of AST and ALT enzymes that are abnormal.

### • The most common HCV genotype

To determine the circulating HCV isolates, seropositive samples were genotyped. Out of 76 patients, 34.2% were genotype 1, 2.6% genotype 2, 14.5% genotype 3 and the highest of all 48.6% were genotype 4 (Table 2). In this study, HCV genotype 4 was the major genotype at least among this group of Saudi Arabian patients.

**Table 2.**
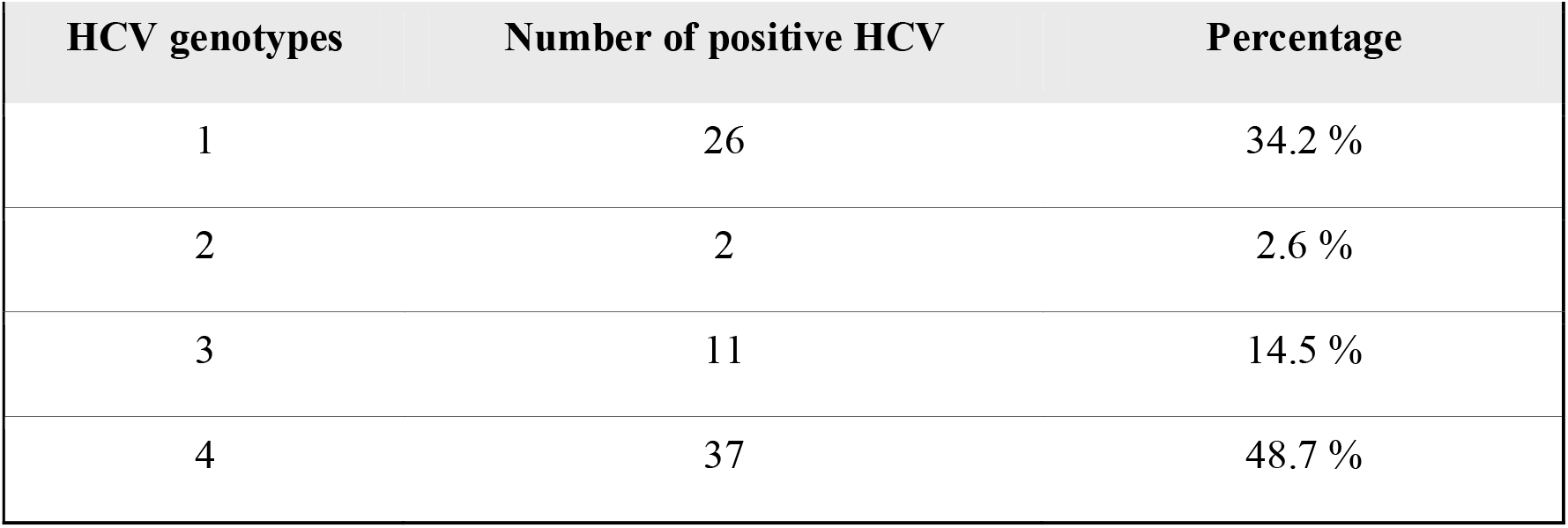
Percentage of HCV genotypes (n=76).

### • Age, gender and genotype

Patients’ sex and age were considered in the analysis of HCV genotypes. We then analyzed gender-related differences observed between the genotypes as shown in Figure 2 and Table 3. Genotypes 1, 3 and 4 were higher in females than males, possibly due to a higher prevalence of blood transfusion among women than men. The association between age and genotype is rather more interesting. In order to investigate the relationship between age and genotypes, Figure 3 and Table 4 summarize the overall association between age group and all tested HCV genotypes. Two male patients were genotype 2 positive; 70 and 75 years old. The rest of genotypes were found in young as well as elderly patients (Figure 3 and Table 4). However, genotype 3 was only seen in patients in their thirties and fifties.

**Figure. 2.**
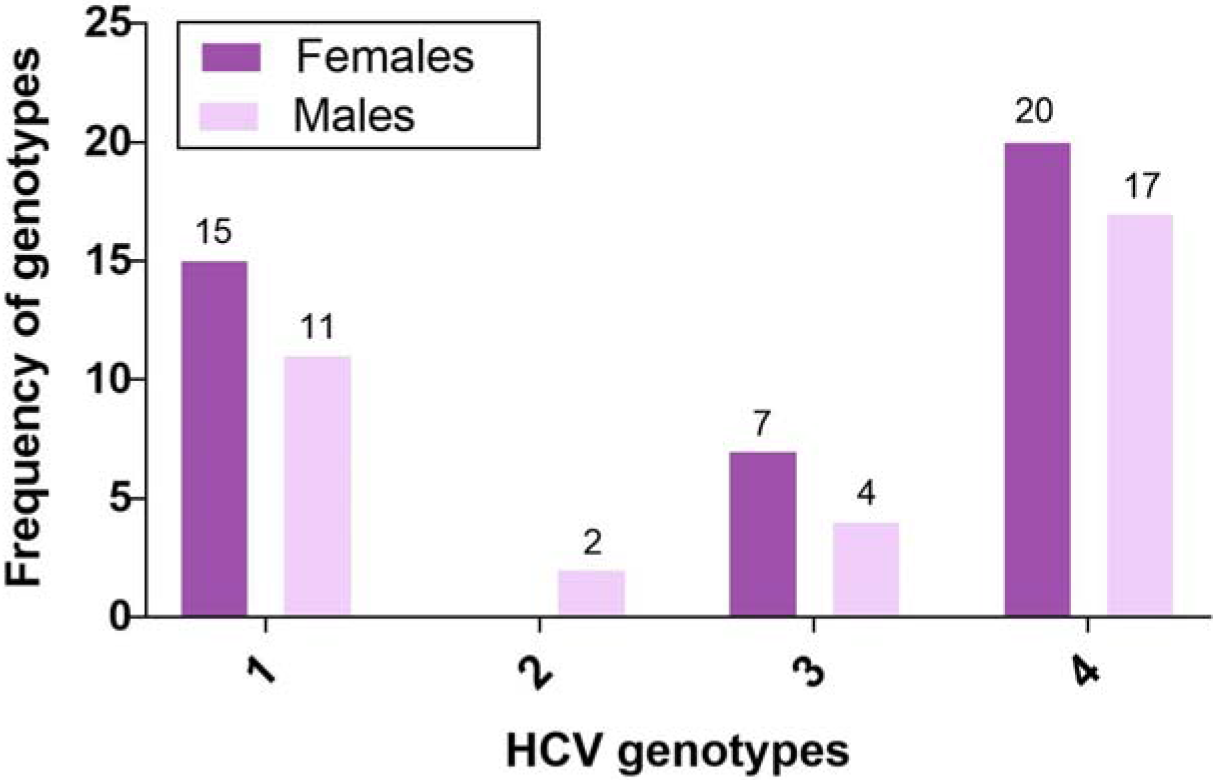
Sex versus HCV genotypes. The frequency of HCV genotypes with sex. Figure shows the detailed values of the rate with the total and percentage. M: Male; F: Female.

**Figure 3.**
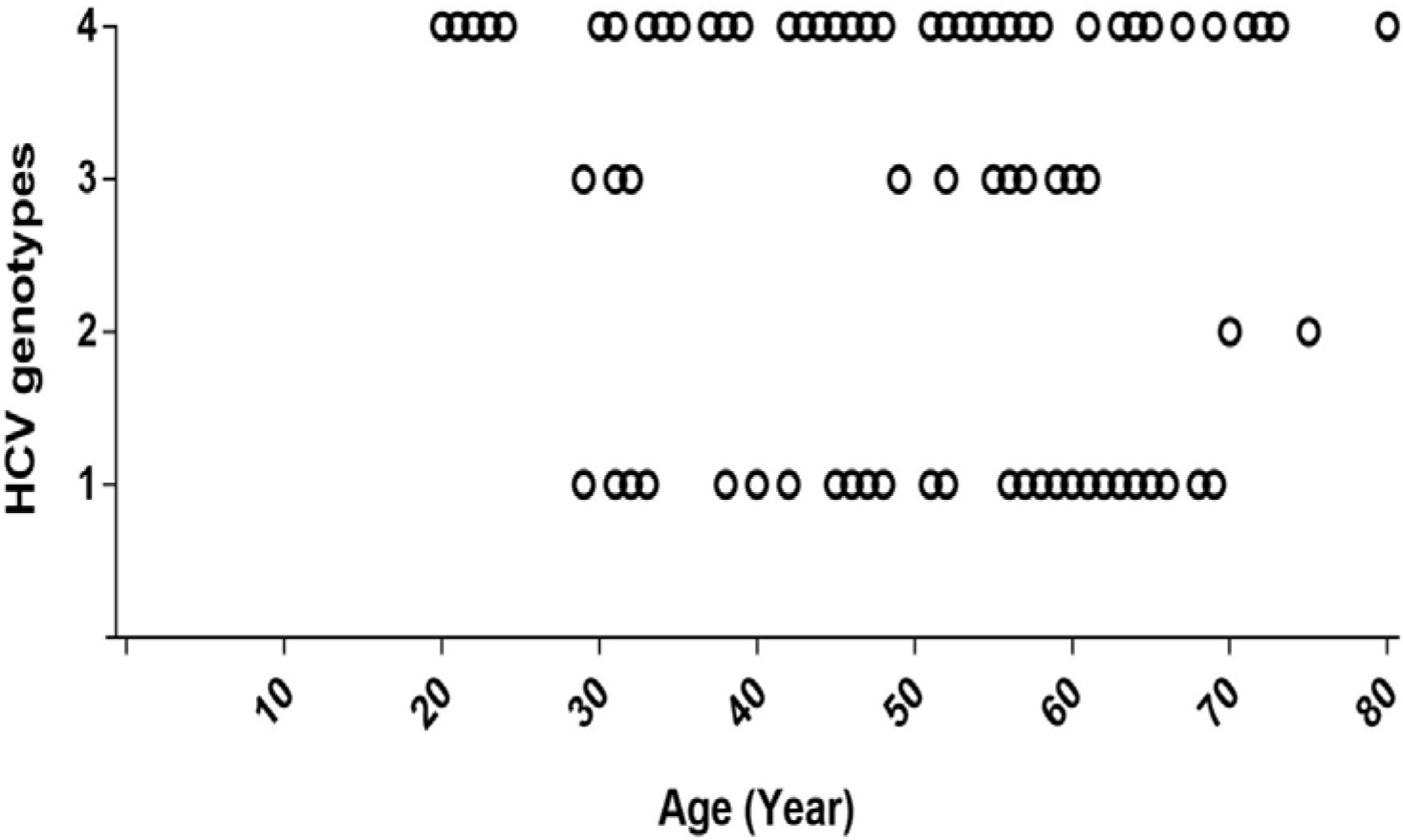
Age versus genotype. Genotypes (blue circles) were aligned with the age.

**Table 3.**
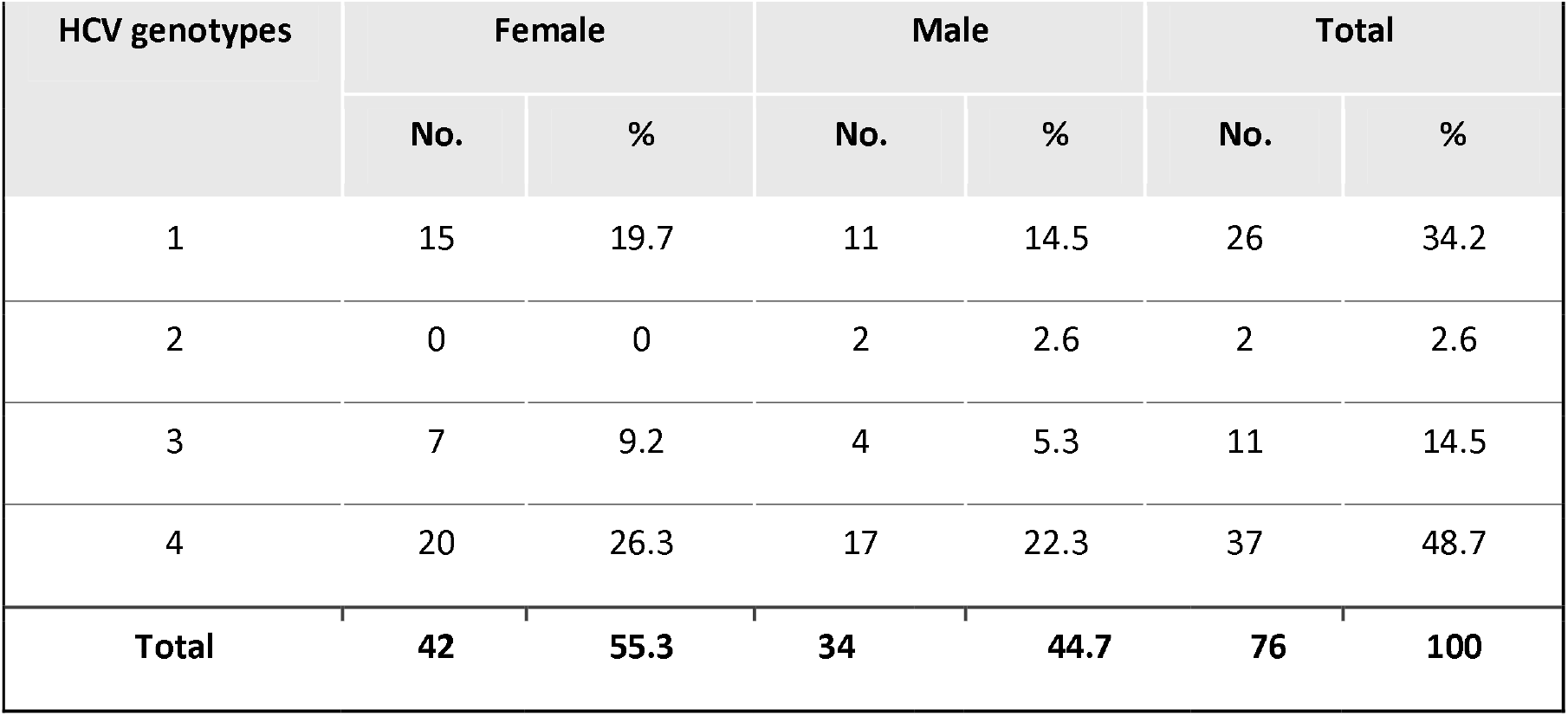
HCV genotypes variation among Saudi Arabian Females and Males.

**Table 4.** HCV genotypes distribution in different age group.

| Age group | Female |  | Male |  | Total |
| --- | --- | --- | --- | --- | --- |
|  | No. | % | No. | % |  |
| 0-20 y | 0 | 0 | 1 | 100 | 1 |
| 20-40 y | 10 | 45 | 12 | 55 | 22 |
| 40-60 y | 24 | 71 | 10 | 29 | 34 |
| 60-80 y | 8 | 42 | 11 | 58 | 19 |
| **Total** | **42** | **55** | **34** | **45** | **76** |

**Table 5.** Frequency of HCV genotypes and history of blood transfusion.

| History of blood transfusion | HCV genotypes |  |  |  | Total / percentage |
| --- | --- | --- | --- | --- | --- |
|  | 1 | 2 | 3 | 4 |  |
| Yes | 7 | 1 | 1 | 4 | 13 (17.1 %) |
| No | 19 | 1 | 10 | 33 | 63 (82.9 %) |

### • Blood donation and HCV infections

Sixty-three patients (82.9%) were non-transfusion recipients, while the rest (13 patients) have been subjected to blood transfusion in the past. All four genotypes found in this investigation were considered in the analysis of blood transfusion. Genotype 4 was highest among non-transfusion patients. However, genotype 1 was the most prevalent among blood transfusion recipients (Figure 4).

**Figure 4.**
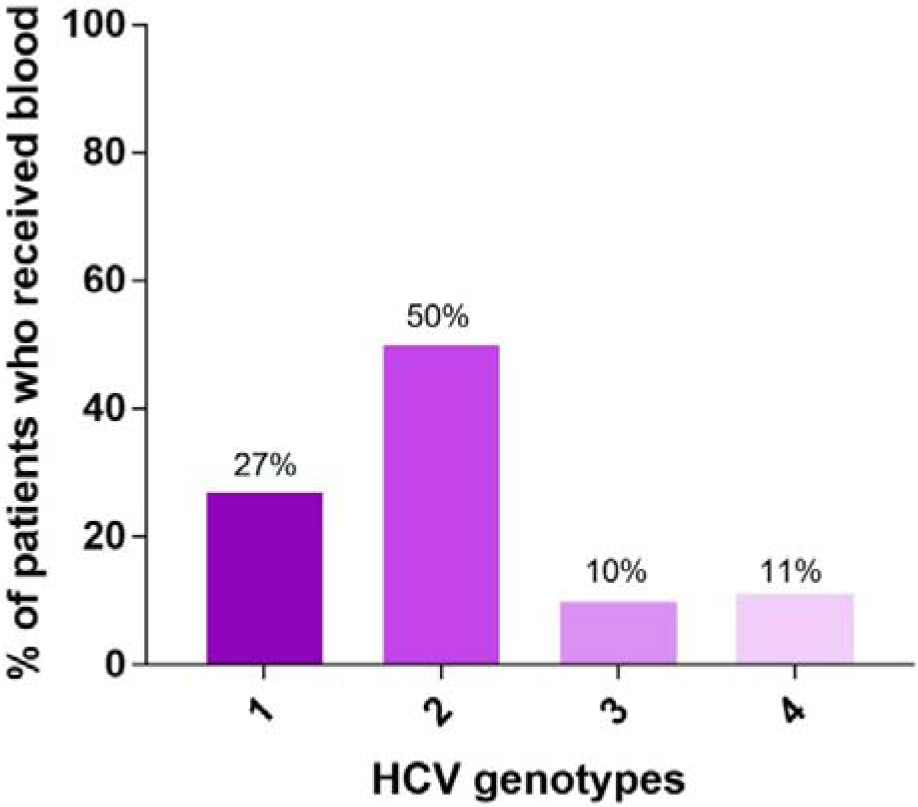
Frequency of HCV genotype and history of blood transfusion.

## Discussion

Data on the prevalence of circulating HCV strains in the Southern Region, Saudi Arabia, is limited. Determination of HCV genotypes has become crucial for the the assessment of clinical course and administration of antiviral drugs. Our investigation attempted to give insight towards the actual genotypes present in the region and to assess the varied epidemiology of the genotypes. HCV chronic patients were followed up by determining the viral load quantitatively. Samples with no viral load determined were either acute or from different clinics (Table 1).

More than one-third of patients were asymptomatic with normal liver enzymes. Both ALT and AST parameters (Table 1) showed significant differences between HCV genotypes. Many studies have shown that HCV genotype 1 is linked with severe liver disease and elevated liver enzymes [27]. It has been estimated that around 25% of chronic hepatitis C have normal ALT level [28]. However, the level of ALT is known to be fluctuating in patients with chronic hepatitis C and flares of disease activity [29].

Al-Raddadi *et. al*. 2017 claimed that the prevalence of HCV in Saudi Arabia is among the lowest worldwide, but the infection rate found in this study suggests that the prevalence could be higher than expected. A previous study showed that 5.87% of Saudis were seropositive to HCV [30]. Moreover, it has been estimated that 21.3 million individuals in the Eastern Mediterranean countries are HCV carriers, similar to Europe and the Americas combined [31]. The rate of HCV infection among the Injecting Drug Users (IDUs) in Saudi Arabia is relatively high [32], and this demands preventive strategies for improved socioeconomic welfare.

A recent study in the Western Province showed that HCV genotype 1 is the most predominant [33]. In this study, we determined that HCV genotype 4 was the most common (48.7%) in the Southern Province. This finding is consistent with other studies [18–20,30,34] in which genotype 4 is widespread among the population. Moreover, a previous investigation detected that 50% of Saudi Arabian patients were infected with genotype 4 [35]. However, their sample size (n=22) was less than the number of patients used in this study (n=76).

Among hundreds of patients visited AFHSR clinics during collection of samples for HCV assays, we found that the rate of infection in females is higher than males (55.3% vs. 44.7% respectively). Bawazir *et. al*. 2017 surveyed 630 HCV-infected Saudi patients, and found that 47.6% were males and 52.4% were females. Discrepancies exist in results regarding gender and pathogenesis [36]. This requires further evaluation to draw evidence-based conclusions. In this study, no association was found between sex and genotype. In consistency with our investigation, Al Traif *et al*. 2013 study has not found a relationship between gender and genotype.

Interestingly, we found that the least prevalent genotype was exclusive to elderly patients. Genotype 2 was only seen in patients above 70yrs old, despite the small sample size with this particular genotype. In addition, Farag *et. al*. 2015 investigated 20 Egyptians and 20 Saudi Arabian patients. They observed that genotype 2 was only detected in Saudi patients.

Patients in dialysis settings are vulnerable to HCV infection despite good hygienic measures [37]. This necessitates strict control precautions and preventive methods to minimize infection [38]. Here, we focused on the history of blood transfusion. The relationship between blood transfusion and the most common genotypes (gt 4 and gt 1 respectively) were examined. An interesting observation is that patients with history of blood transfusion showed a high percentage of genotype 1 compared to the total number of all patients with genotype 1. The percentage is more than double with genotype 1 than genotype 4 (23% and 11% respectively). One limitation is that the overall number of patients infected with genotype 4 is higher than genotype 1. Whether this is a coincidental observation or with a significant link, it is worth noting. Another surprising observation is that the first five patients with the highest viral load have all received blood transfusion. It is difficult to infer any conclusion with regards to these unexpected observations found with the transfused patients, but it is noteworthy to consider in future investigations.

## Conclusion

Out of 76 HCV-positive patients, genotype 4 was the highest among all assessed genotypes. This result was consistent with other findings. No association was found between gender, age and genotype. Genotype 2 was only found in elderly patients aged 70 and 75 years old. Genotype 1 was common among transfused patients.

## Future perspective

Future studies will focus on subtypes and phylogenetic studies with higher number of samples as well as to determine whether subtype 4d is more common than other subtypes or not [39]. The investigation will include the effect of low oxygen level on the severity of HCV infection [40] hence these patients live at high altitude. This will determine any interesting or rare mutations.

## Financial and competing interests disclosure

The authors have no relevant affiliations or financial involvements with any organization or any financial interest with the subject matter or materials discussed in the manuscript. This includes employment, consultancies, honoraria, stock ownership or options, expert testimony, grants or patents received or pending or royalties.

No writing assistance was utilized in the production of this manuscript.

## Ethical conduct of research

The authors state that they obtained appropriate institutional review board approval or have followed the principles outlined in the Declaration of Helsinki for all human or animal experimental investigations. In addition, for investigations involving human subjects, informed consent has been obtained from the participants involved.

## Executive summary

### The most common HCV genotype

- Genotype 4 was the most prevalent of all assessed HCV genotypes among 76 positive patients.

### Gender and genotype

- No association was found between sex and genotype.

### Association between blood donation and genotype 1

- The increased percentage of genotype 1 among those who have received blood donation in the past is interesting.

